# Last-come, best served? Mosquito biting order and *Plasmodium* transmission

**DOI:** 10.1101/2020.04.08.031625

**Authors:** J. Isaïa, A. Rivero, O. Glaizot, P. Christe, R. Pigeault

## Abstract

A pervasive characteristic of malaria parasite infection in mosquito vector populations is their tendency to be overdispersed. Understanding the mechanisms underlying the overdispersed distribution of parasites is of key importance as it may drastically impact the transmission dynamics of the pathogen. The small fraction of heavily infected individuals might serve as superspreaders and cause a disproportionate number of subsequent infections. Although multiple factors ranging from environmental stochasticity to inter-individual heterogeneity may explain parasite overdispersion, *Plasmodium* infection has also been observed to be highly overdispersed in inbred mosquito population maintained under standardized laboratory conditions, suggesting that other mechanisms may be at play. Here, we show that the aggregated distribution of *Plasmodium* within mosquito vectors is partially explained by a temporal heterogeneity in parasite infectivity triggered by the bites of blood-feeding mosquitoes. Several experimental blocks carried out with three different *Plasmodium* isolates have consistently shown that the transmission of the parasite increases progressively with the order of mosquito bites. Surprisingly the increase in transmission is not associated with an increase in *Plasmodium* replication rate or higher investment in the production of the transmissible stage (gametocyte). Adjustment of the physiological state of the gametocytes could be, however, an adaptive strategy to respond promptly to mosquito bites. Overall our data show that malaria parasite appears to be able to respond to the bites of mosquitoes to increase its own transmission at a much faster pace than initially thought (hours rather than days). Further work needs to be carried out to elucidate whether these two strategies are complementary and, particularly, what are their respective underlaying mechanisms. Understanding the processes underlying the temporal fluctuations in *Plasmodium* infectivity throughout vertebrate host-to-mosquito transmission is essential and could lead to the development of new approaches to control malaria transmission.

**Author summary:** *Plasmodium* parasites are known for being the etiological agents of malaria and for the devastating effects they cause on human populations. A pervasive characteristic of *Plasmodium* infection is their tendency to be overdispersed in mosquito vector populations: the majority of mosquitoes tend to harbour few or no parasites while a few individuals harbor the vast majority of the parasite population. Understanding the mechanisms underlying *Plasmodium* overdispersed distribution is of key importance as it may drastically impact the transmission dynamics of the pathogen. Here, we show that the aggregated distribution of *Plasmodium* parasites within mosquito vectors is partially explained by a temporal heterogeneity in *Plasmodium* infectivity triggered by the bites of blood-feeding mosquitoes. In other words, mosquitoes that bite at the beginning of a 3h feeding session have significantly fewer parasites than those that bite towards the end. Malaria parasite is therefore capable of responding to the bites of mosquitoes to increase its own transmission at a much faster pace than thought (hours rather than days). Understanding the processes underlying the temporal fluctuations in *Plasmodium* infectivity throughout vertebrate host-to-mosquito transmission is essential and could lead to the development of new approaches to control malaria transmission.

## Introduction

An ubiquitous feature of parasite infections is their tendency to be overdispersed or aggregated [1–4]. In other words, in a natural population of hosts, the majority of individuals tend to harbour few or no parasites while a few hosts harbour the vast majority of the parasite population. This pattern has been observed in a wide range of diseases ranging from viruses and fungal parasites of plants [5,6] to protozoan and metazoan parasites of humans [7,8].

Previous work has shown that the overdispersed pattern of parasites among hosts can have strong repercussions for disease dynamics [9,10]. Overdispersion reduces the deleterious effects of parasites on host populations but also increases the intensity of density-dependent suppression of parasite population growth (e.g. mating probability, intra- and inter-specific competition [11,12]). Another property emerging from parasite overdispersion is the effect on infectious disease epidemiology and parasite transmission. The small fraction of heavily infected individuals may serve as super-spreaders and therefore play a large role in disease transmission [13–15]. In many host–parasite systems, 20% of hosts are responsible for 80% of new infections [16,17]. In vector-borne diseases, parasite overdispersion has been observed both in the vertebrate host and in the vector populations [18–22]. Despite this, studies to date have mainly focused on the epidemiological consequences of parasite overdispersion in vertebrate host rather than vectors [17,23,24]. Yet, for many of these diseases, key traits determining the transmission dynamics of the pathogen such as the lifespan and feeding behaviour of vectors as well as the length of the parasite’s extrinsic incubation period may depend on the intensity of parasite infection in the vector [25–29].

Anderson and Gordon (1982) identified environmental stochasticity as the prime cause of overdispersion in parasite populations [30]. This included not only the physical parameters of the environment, but also the differences in host susceptibility to the infection induced by behavioural differences, genetic factors or varying past experiences of infection. The mechanisms underlying the aggregated distribution of parasites in vector populations remains however rarely explored and little understood.

*Plasmodium* parasites are known for being the etiological agents of malaria and for the devastating effects they cause on human populations in the African continent. These vector-borne parasites are however also found infecting many other terrestrial vertebrate species, including other mammals, reptiles and birds. The life cycle of the parasite is the same in all hosts, irrespective of their taxa. When the mosquito vectors take a blood meal on an infected host, they ingest the parasite’s transmissible stages (female and male gametocytes). After the sexual reproduction of the parasite, the motile zygotes penetrate the wall of the midgut and start developing into oocysts, which in turn produce the transmissible sporozoites in the mosquito’s salivary glands. There is abundant evidence showing that the distribution of oocysts, the most commonly quantified parasite stage in mosquitoes, is highly overdispersed [7,31–34]. The most straightforward explanation for this aggregated distribution of oocysts is that it is driven by heterogeneity in vector susceptibility to *Plasmodium* infection associated to their genetic background or to their physiological status [7,35,36]. Polymorphism in mosquito immune genes is strongly associated with natural resistance to *Plasmodium* [35,37] and aging also tends to decrease the susceptibility of vectors to *Plasmodium* infection [36]. Puzzlingly, however, oocyst overdispersion is also extremely common under controlled laboratory conditions in highly inbred, and therefore physiologically and genetically homogeneous, mosquito populations [7,31,33,34]. This suggests that genetic or physiological heterogeneities between mosquitoes may only be part of the explanation, and that the causes of the aggregated distribution of oocysts in vectors may also lie elsewhere.

One possible explanation is the existence of *spatial* aggregation of gametocytes in the vertebrate blood. Recent work has shown that gametocyte densities in humans can vary in as much as 45% in blood collected from different parts of the body (Pigeault et al *.in prep).* Although the direct connection between spatial heterogeneity in blood and overdispersion in mosquitoes has never been made, it has been reported that *Plasmodium* gametocytes show an aggregated distribution within mosquitoes which recently fed on human host [38].

Alternatively, the aggregated distribution of *Plasmodium* parasites within mosquitoes could be due to a within-host *temporal* variation in parasite densities and/or infectivity. Under this scenario, mosquitoes feeding during the high parasite density/infectivity phase would be more heavily infected than those feeding during the low density/infectivity phase*. Plasmodium* parasite density and/or infectivity in the vertebrate host can indeed vary within relatively short temporal intervals. A recent study found that rodent malaria *P. chabaudi* gametocytes are twice as infective at night despite being less numerous in the blood [39]. A periodic late afternoon increase in parasitemia is also observed in the avian malaria system [40]. Such temporal variations may respond to changes in the physiological, nutritional or immunological condition of the host [41–43]. They may, however, also be an adaptive strategy of the parasite aimed at maximizing its own transmission [40,44]. Recent work has shown that host parasitaemia increases a few days after a mosquito blood feeding bout, suggesting that *Plasmodium* may be capable of adjusting its transmission strategy by responding plastically to the temporal fluctuations in vector availability [40,44]. These results, however, are not able to explain the aggregated distribution of parasites among mosquitoes feeding within a short feeding bout typically lasting a handful of hours.

Here, we test whether the *Plasmodium* is able to respond plastically to the bites of mosquitoes at a much more rapid pace than initially thought. More specifically, we test whether there is a pattern in the oocyst load of mosquitoes feeding within a short (3-hour) time interval: do the first mosquitoes to bite the host increase the infectivity of the parasite so that mosquitoes biting last end up with significantly increased oocystaemias? To test this hypothesis, we use the avian malaria experimental system, the only currently available animal experimental system that allows working with a parasite recently isolated from the wild (*Plasmodium relictum*), with its natural mosquito vector (the mosquito *Culex pipiens*). Specifically, we carry out a series of experiments designed to answer the following two main questions: 1) Is oocystaemia correlated with mosquito biting order? In other words, do mosquitoes biting first have a lower intensity of infection than those biting later on? and 2) Is this due to a temporal increase in the parasitaemia/gametocytaemia of birds as a result of mosquito bites?

## Results

To investigate the impact of mosquito bite-driven plasticity on *Plasmodium* transmission we used the avian malaria biological system [45]. Birds (*Serinus canaria*) infected by *Plasmodium relictum*, the causative agent of the most prevalent form of avian malaria in Europe, were exposed to a wild-caught lineage of *Culex pipiens* mosquitoes for 3 hours (6 – 9 p.m.). Mosquitoes were sampled at regular intervals thereafter (different protocols for the three experiments, see below) and dissected one week later to count the number of oocysts in the midgut. To investigate the impact of vector bites on parasite population growth, the parasitaemia (number of parasites in the blood) and gametocytaemia (number of gametocytes in the blood) of vertebrate hosts exposed or not (control) to mosquitoes were measured just before and just after the mosquito exposure period (6 – 9 p.m.).

### Experiment 1: Oocyst burden and mosquito biting order: batch experiment

In the first experiment, birds were exposed to four successive batches of 25 ± 3 uninfected mosquitoes. Each mosquito batch was kept in the cage for 45 minutes before being replaced with a new batch (batch 1 (T0_min_), batch 2 (T45_min_), batch 3 (T90_min_) and batch 4 (T135_min_)). At the end of each exposure period, all mosquitoes were removed from the cages and were immediately replaced by a new batch. The blood meal rate (*i.e.* proportion of blood-fed mosquitoes) and the haematin quantity, a proxy for blood meal size, were similar for all batches (model 1: χ² = 5.90, p = 0.116, model 2: χ²= 3.55, p = 0.314 respectively, statistical models are described in **Table S1**). Although mosquitoes from batches 3 and 4 tend to have a higher infection prevalence (proportion of mosquitoes containing at least 1 oocyst in the midgut; mean ± SE: batch 3: 64.4% ± 11.9, batch 4: 78.2% ± 8.6) than those from batches 1 and 2 (batch 1: 56.7% ± 15, batch 2: 56.7% ± 19.4) the difference in prevalence between the different batches was not statistically significant (model 3: χ² = 2.74, p = 0.433). The overall distribution of oocyst burden across batches was highly overdispersed (Figure 1A, mean ± se Variance-to-Mean Ratio = 11.48 ± 3.37). Oocyst burden increased with mosquito batch (geometric mean: batch 1: 3.41 ± 3.04, batch 2: 3.99 ± 3.25, batch 3: 6.13 ± 3.36 and batch 4: 11.84 ± 3.53, model 4: χ² = 35.283, p < 0.0001, Figure 1A). Females from batch 4 had almost twice as many oocysts as those from batch 3 (contrast analyses: batch4/batch3: χ² = 11.02, p < 0.001) and three times more than females from batches 1 and 2 (batch4/batch2: χ² = 17.95, p < 0.001, batch4/batch1: χ² =19.31, p < 0.0001, Figure 1A). No significant difference was however observed between mosquitoes from batches 1 and 2 (contrast analyses: batch1/batch2: χ² = 0.15, p = 0.697) or between mosquitoes from batches 2 and 3 (batch2/batch3: χ² = 2.29, p = 0.129, Figure 1A). Haematin quantity had no effect on the oocyst burden (model 4: χ² = 0.02, p = 0.875).

The increase in *Plasmodium* oocyst burden with mosquito batch was not explained by an increase in total parasite or gametocyte burden in the birds’ peripheral blood. The parasitaemia and gametocytaemia of exposed birds remained roughly constant between the beginning and the end of the experiment (parasitaemia= model 5: χ² = 0.39, p = 0.529, gametocytaemia = model 6: χ² = 0.02, p = 0.877 respectively) and were similar between exposed and control (unexposed) birds (parasitaemia = model 5: χ² = 0.29, p = 0.5907, gametocytaemia = model 6: χ² = 0.60, p = 0.4364).

To test the repeatability of our results, a second experimental block, with a new *Plasmodium relictum* strain freshly collected in the field from an infected House sparrow (*Passer domesticus*), was performed. The results of block 2 fully confirmed those of the first block. The blood meal rate and the quantity of haematin excreted by mosquitoes was similar for all batches (model 7: χ²= 1.77 p = 0.621, model 8: χ²= 1.13, p = 0.770). The difference in infection prevalence between the different batches was not statistically significant (model 9: χ² = 5.70, p = 0.127) although mosquitoes from batches 2, 3 and 4 tend to have a higher prevalence (mean ± SE, batch 2: 73.1 % ± 7.0, batch 3: 69.4% ± 7.8 and batch 4: 70.3% ± 7.6 %) than those from batch 1 (mean ± SE, batch 1: 51.1% ± 7.5). The distribution of oocyst burden in mosquitoes was overdispersed (Figure 1B, mean ± se VMR = 11.40 ± 5.66) and we observed a significant increase in oocyst burden with mosquito batches order (model 10: χ² = 30.73, p < 0.0001, geometric mean: batch 1: 3.47 ± 2.69, batch 2: 3.52 ± 2.95, batch 3: 6.06± 2.85 and batch 4: 10.38 ± 2.76, all contrast analyses were significant, Figure 1B). A significant positive correlation between haematin and oocyst burden was found (model 10: χ² = 4.46, p = 0.03). As in the previous experimental block, the vertebrate host parasitaemia and gametocytaemia remained constant between the beginning and the end of the experiment (parasitaemia = model 11: χ² = 1.29, p = 0.256, gametocytaemia = model 12: χ² = 0.88, p = 0.349 respectively) and were similar between exposed and control (unexposed) birds (parasitaemia= model 11: χ² = 2.44, p = 0.118, gametocytaemia = model 12: χ² = 2.45, p = 0.117 respectively).

**Figure 1:**
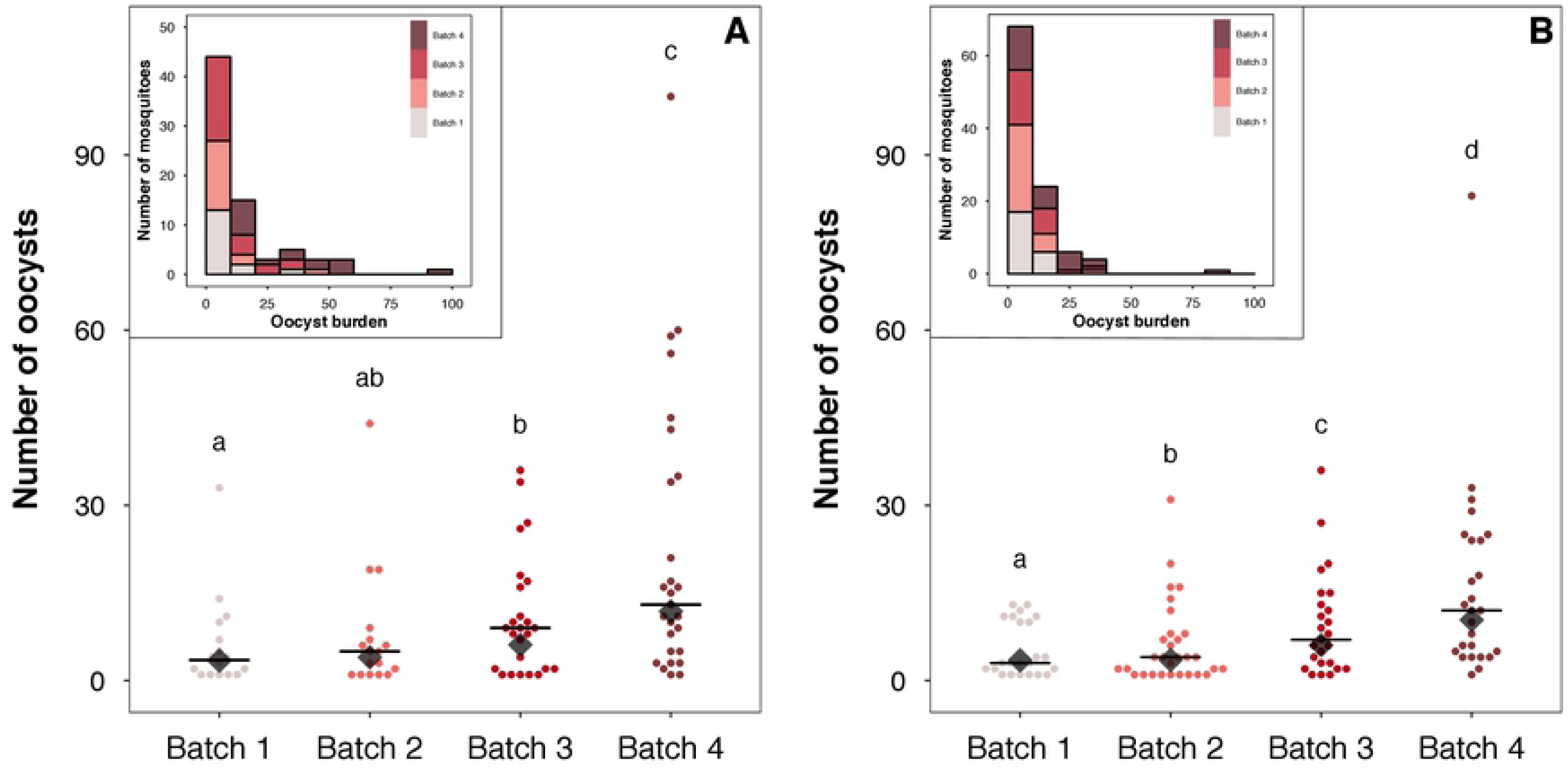
Experiment 1: Impact of mosquito batch order on *Plasmodium* transmission. Number of oocysts in the midgut of *Plasmodium-*infected mosquitoes according to mosquito batches. Each mosquito batch was left for 45 minutes in contact with birds (batch 1 (T0_min_), batch 2 (T45_min_), batch 3 (T90_min_) and batch 4 (T135_min_)). Birds were either infected by a *Plasmodium relictum* lab strain (experimental block 1, panel **A**) or by a *Plasmodium relictum* strain freshly collected in the field (experimental block 2, panel **B**). Black horizontal lines represent medians and black diamond represent geometric means. Levels not connected by same letter are significantly different. Histograms in each panel show the distribution of oocyst burden in mosquitoes in the experimental blocks 1 (**A**) and 2 (**B**), the colors represent the mosquito batches (from 1 to 4).

### Experiment 2: Oocyst burden and mosquito biting order: individual monitoring

A second experiment, with another *Plasmodium relictum* strain freshly collected in the field from an infected Great tit (*Parus major*), was carried out to obtain a finer measurement of the impact of mosquito biting order on their oocyst burden. Here, infected individuals were exposed to 100 mosquitoes for 3h (6.00 pm – 9.00 pm). Cages were continuously observed and mosquitoes were individually removed from the cages immediately after their blood feeding bout. The order of biting of each individual female was recorded.

Haematin quantity and infection prevalence were independent of the mosquito biting order (model 13: χ² = 2.44, p = 0.118, model 14: χ² = 0.83, p = 0.363, respectively). The distribution of oocyst burdens across all mosquitoes was highly overdispersed (mean ± SE. VMR = 90.26 ± 41.53, Figure 2A). Biting order was a significant explanatory factor of oocyst burden: mosquitoes that bit later showed higher oocyst burden than mosquitoes that bit first (model 15: χ² = 8.28 p = 0.004, Figure 2A). A significant positive correlation between haematin quantity and oocyst burden was found (model 15: χ² = 19.151, p < 0.001). As for the first experiment, vertebrate host parasitaemia and gametocytaemia remained constant between the beginning and the end of the experiment (parasitaemia = model 16: χ² = 2.03, p = 0.154, gametocytaemia = model 17: χ² =0.13, p = 0.718 respectively) and were similar between exposed and unexposed (control) birds (parasitaemia = model 16: χ² = 0.98, p = 0.321, gametocytaemia = model 17: χ² = 0.12, p = 0.731 respectively).

**Figure 2:**
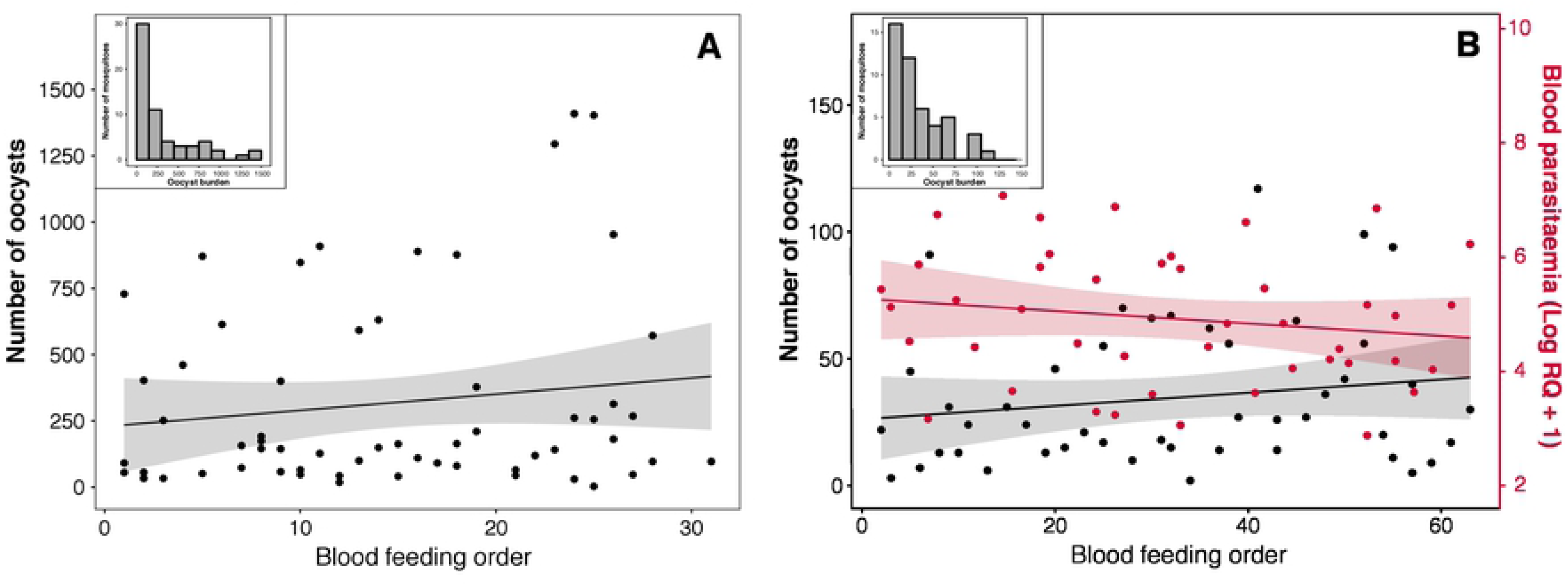
Effect of individual mosquito blood feeding order on the number of parasites ingested and on the intensity of infection. **(A)** Relationship between oocyst burden and mosquito biting order (experiment 2). **(B)** Relationship between the number of parasites ingested (Log(RQ+1), in red), or the oocyst burden (in black), and the mosquito biting order (experiment 3). Each point represents one blood-fed mosquito. Shaded areas on either side of the regression line represent 95% confidence intervals. Histogram in each panel show the distribution of oocyst burden in mosquitoes in the experiment 2 (**A**) and 3 (**B**).

### Experiment 3: Number of parasites ingested and mosquito biting order

The first two experiments showed an increase in the oocyst burden with the order of mosquito bites without, however, showing an increase of the parasite density in the peripheral blood of vertebrate hosts (measured from blood samples). We carried out a third experiment to determine whether the total number of parasites in the blood meal, immediately after the blood feeding, fluctuated during the feeding bout. As for the experiment 2, birds were exposed to 100 mosquitoes for 3h (6.00 pm – 9.00 pm) and mosquitoes were individually removed from the cages immediately after blood feeding. Every second mosquito collected was either immersed immediately in liquid nitrogen or stored in plastic tubes and dissected one week later to count the number of oocysts in the midgut. Frozen blood-fed mosquitoes were used to quantify the number of parasites ingested by qPCR.

The amount of parasite ingested by the mosquitoes remained roughly constant throughout the exposure period (model 18: χ² = 1.54, p = 0.215 Figure 2B). The hematin quantity and the infection prevalence (oocyst stage) were also independent of the mosquito biting order (model 19: χ² = 1.89, p = 0.169, and model 20: χ² = 0.37, p = 0.545 respectively). In contrast, the distribution of oocyst burden across all mosquitoes was still overdispersed (mean ± SE. VMR = 15.03 ± 1.86, Figure 2B) and was significantly explained by the mosquito biting order (model 21: χ² = 6.45, p = 0.011, Figure 2B). As in the experiment 2, mosquitoes that bit later showed higher oocyst burden than mosquitoes that bit first (Figure 2B).

## Discussion

Overdispersed distribution of vector-borne parasite within vertebrate and invertebrate host populations has profound consequences on parasite transmission and disease control strategies [16,28,46]. Parasite overdispersion is driven by multiple factors ranging from population processes to inter-individual heterogeneity in susceptibility and parasite exposure [11,47–49]. Here, using three different isolates of *Plasmodium relictum*, we provide evidence that the aggregated distribution of malaria parasites within mosquito vectors may also be explained by the mosquito biting order: mosquitoes that bite first have a lower intensity of infection than those that bite later on. This fluctuation in *Plasmodium* infectivity may reflect an adaptive strategy of parasites selected to optimize transmission.

The abundance of invertebrate vectors fluctuates at time scales ranging from daily to annual [40,50–52]. Previous studies have shown that malaria parasites have evolved two different and complementary transmission strategies to cope with both short (circadian) and long (seasonal) term fluctuations in mosquito activity. *Plasmodium* adopts an unconditional strategy whereby within-host parasitaemia and/or gametocyte infectivity increases daily when its vector is most active [39,40] but also a plastic strategy allowing parasite growth to increase after exposure to mosquito bites [40,44,53]. This plastic strategy allows the parasite to react to daily and seasonal fluctuations in mosquito abundance [40,44].

In this study we demonstrate that *Plasmodium* plastic response is much faster than initially thought [40,44]. When vertebrate hosts were exposed to mosquito bites during a short period of time (3 hours), parasite transmission increased gradually with the biting order of mosquitoes. *Plasmodium* transmission was tripled between the first and the last blood fed mosquito. Although the biting order of the mosquito cannot be decoupled from the biting time (these two parameters are obviously highly correlated), the increase in transmission in such a short period of time suggests that the effect observed here cannot be explained solely by circadian fluctuation in parasite density in vertebrate blood. Many mosquito species exhibit a circadian rhythm in the host-biting activity [40,50] but stochastic environmental factors such as variations in temperature, wind or humidity impact drastically the abundance of mosquitoes from one day to another [54–56]. Therefore, the association between an unconditional strategy (circadian fluctuation) and a quick plastic response to mosquito bites may allow malaria parasites to fine-tune investment in transmission according to the presence of mosquitoes.

Interestingly, this adaptive hypothesis involving an active parasite response to mosquito bites is not mediated by an increase in either parasite replication rate or gametocyte production: parasitaemia and gametocytaemia of birds exposed to mosquitoes were not different before and after mosquito probing. This result was confirmed by monitoring the number of parasites ingested by the mosquitoes immediately after the blood meal, throughout the exposure period. These results contrast with those obtained in recent studies [40,44] where the increase in oocyst burden observed in mosquitoes fed on a host a few days after the host was exposed to vector bites was correlated with an increase in parasitaemia and gametocythemia. Our study suggests that malaria parasite have evolved an alternative strategy acting at a shorter term. This strategy could be to adjust the physiological state of the gametocytes to respond promptly to mosquito bites. It has been suggested as early as 1966 [57] that malaria parasite infectivity is not only due to the number of gametocytes in the blood but also to their physiological state. This prediction was recently experimentally confirmed by a study carried out on rodent malaria parasite: *P. chabaudi* gametocytes were twice as infective at night despite being less numerous in the blood [39]. Mechanisms underlying gametogenesis remains poorly understood. Although we know that gametocytes go through several stages of development before reaching the so-called “mature” stage, from 1 to 8 stages depending on the species of *Plasmodium* [58], we do not know whether mature stage is systematically infectious. The ability of malaria parasites to accelerate the rate of maturation and/or infectivity of gametocytes in response to mosquito bites should be explored.

Alternatively, the response of the vertebrate host to mosquito bites could also enhance parasite transmission from the vertebrate host to the invertebrate vector by two non-exclusive mechanisms: (**i**) increased infectivity and/or survival of parasites in vector midgut and (**ii**) modified susceptibility of mosquitoes to infection. *Plasmodium* abundance experiences strong fluctuations during its journey within the mosquito, which are partly intertwined with the kinetics of blood digestion [32]. Within seconds of ingestion into the mosquito blood meal, the drop in temperature and the rise in pH, associated to the presence of xanthurenic acid, triggers gametocyte activation and differentiation into gametes [59–61]. Studies on ookinete production have revealed that not only mosquito-derived xanthurenic acid but also undefined blood-derived factors ingested by mosquito are significant sources of gametocyte activating factor [62,63]. Indeed, numerous host blood-derived compounds remain or become active through the mosquito blood digestion, especially since the parasite is no longer protected by the red blood cell membrane. For instance, complement components, vertebrate antibodies or regulator factor H, may impact gametocytes-to-zygote and zygote-to-ookinetes stages transition and survival [64–66]. Several studies also showed that ingested vertebrate-derived factors negatively impact mosquito microbiota (e.g. complement cascade [67,68]) and their peritrophic matrix (e.g. chitinase [66,69]) both known to play a key role in the mosquito refractoriness to *Plasmodium* infection [70,71]. The concentration of these vertebrate-derived compounds in the ingested blood and, ultimately, their impact on parasite infectivity and/or vector susceptibility, might vary progressively as the number of bites increases and thus explain the increase in oocyst density with mosquito biting order.

In summary, we provide evidence that the overdispersion of parasite burden observed in mosquitoes fed on a same infected host is partly explained by a temporal heterogeneity in *Plasmodium* infectivity resulting from the biting order of mosquitoes. These results show that the parasite is either directly or indirectly capable of responding to the bites of mosquitoes to increase its own transmission at a much shorter time scales than initially thought (hours rather than days [40,44]). Further work needs to be carried out to elucidate whether these two strategies are complementary and, particularly, what are their respective underlaying mechanisms. According to estimates by the World Health Organization, 228 million cases of human malaria occurred in 2018, with 405 000 resulting in death. Despite recent progress towards disease control, the number of malaria cases has increased in several countries. The efficacy of control strategies is continually challenged and threatened by the evolution of insecticide [72] and drug [73] resistances. To overcome these issues, the development of innovative therapeutic approaches is necessary and urgent. Understanding the mechanisms allowing *Plasmodium* to increase transmission in response to mosquito bites could lead to the development of new pharmaceutical approaches to control malaria transmission.

## Materials and Methods

### Malaria parasites and mosquito vector

*Plasmodium relictum* (lineage SGS1) is the most prevalent form of avian malaria in Europe [74]. The parasite strain used in the first block of the first experiment was isolated from an infected Great tit (*Parus major*) in 2015. The parasite used in the second experiment was isolated from an infected Great tit (*Parus major*) in April 2018. The parasite strain used in the second block of the first experiment and in the third experiment was isolated in January 2019 from an infected House sparrow (*Passer domesticus*). All strains were maintained by carrying out regular passages across our stock canaries (*Serinus canaria*) through intraperitoneal injections (i.p) until the beginning of the experiment.

All the experiments were conducted with a lineage of *Culex pipiens* mosquitoes, the main vector of *Plasmodium relictum* in Europe, collected in Lausanne (Switzerland) and maintained in insectary since August 2017. Mosquitoes were reared using standard protocols [75]. We used females 7-13 days after emergence that had no prior access to blood. Mosquitoes were maintained on glucose solution (10%) since their emergence and were starved (but provided with water to prevent dehydration) for 24h before the experiment.

### Experimental design

Prior to the experiments, a small amount (ca.3-5 µL) of blood was collected from the medial metatarsal vein of each canary to ensure that they were free from any previous haemosporidian infections [76]. Birds were inoculated by intraperitoneal injection of 100µL of an infected blood pool (day 0). The blood pool was made with a 1:1 mixture of PBS and blood sampled from 2-4 canaries infected with the parasite three weeks before the experiment.

#### Experiment 1

The two experimental blocks of the first experiment were carried out with 14 and 5 infected birds respectively. Day 11-13 post-infection, corresponding to the acute phase of infection, blood was sampled from each bird at 5:45 p.m. Straight afterwards blood sampling, birds were placed individually in an experimental cage (L40 x W40 x H40 cm). At 6:00 p.m., 8 and 3 haphazardly chosen birds, for block 1 and 2 respectively, were exposed to mosquito bites following the protocols described below.

Birds from the exposed treatment were exposed to four successive batches of 25 ± 3 uninfected females’ mosquitoes. Each mosquito batch was left in the cage for 45 minutes before being taken out and replaced by a new batch (i.e. batch 1 (T0_min_), batch 2 (T45_min_), batch 3 (T90_min_) and batch 4 (T135_min_)). Blood fed mosquitoes in each batch were counted and individually placed in numbered plastic tubes (30 ml) covered with a mesh with a cotton pad soaked in a 10% glucose solution. At the end of the last mosquito exposure session (9:00 p. m.) a second blood sample was taken from each bird. A red lamp was used to capture blood fed mosquitoes without disturbing the birds and the mosquitoes. Unexposed birds (control) were placed in the same experimental condition but without mosquitoes.

Tubes containing the blood fed mosquitoes were kept in standard insectary conditions to obtain an estimate of the blood meal size and the success of the infection (infection prevalence and oocyst burden). For this purpose, 7-8 days post blood meal, the females were taken out of the tubes and the amount of haematin excreted at the bottom of each tube was quantified as an estimate of the blood meal size [75]. Females were then dissected and the number of *Plasmodium* oocysts in their midgut counted with the aid of a binocular microscope [75].

#### Experiment 2

The same protocol as described above was used for the second experiment, except that birds exposed to mosquitoes (4 of the 8 infected birds) were exposed to a single batch of 100 uninfected mosquitoes for 3h (6:00-9:00 p.m.). Female mosquitoes were continuously observed and individually removed from the cages immediately after blood feeding in order to record the order of biting of each female.

#### Experiment 3

Two infected birds were exposed to a single batch of 100 mosquitoes for 3h (6.00 pm – 9.00 pm) and mosquitoes were removed from the cages immediately after blood feeding. The order of biting of each female was recorded and every second mosquito collected was either immersed immediately in liquid nitrogen to quantify the number of parasites ingested by qPCR or stored in plastic tubes and dissected one week later to count the number of oocysts in the midgut.

### Vertebrate host infection

The parasitaemia (total proportion of red blood cells infected) and gametocytaemia (proportion of red blood cells infected by mature gametocytes, the sexual stages of the parasite) of vertebrate hosts exposed or not (control) to mosquitoes were measured just before and just after the mosquito exposure period (6 – 9 p.m.) using blood smears [74].

### Molecular analyses

The quantification of parasites contained within the blood meal was carried out using a quantitative PCR (qPCR) with a protocol adapted from Cornet et al. (2013). Briefly, DNA was extracted from blood-fed females using standard protocols (Qiagen DNeasy 96 blood and tissue kit). For each individual, we conducted two qPCRs: one targeting the nuclear 18s rDNA gene of *Plasmodium* (Primers 18sPlasm7 5’-AGCCTGAGAAATAGCTACCACATCTA-3’, 18sPlasm8 5’-TGTTATTTCTTGTCACTACCTCTCTTCTTT-3’) and the other targeting the 18s rDNA gene of the bird (Primers 18sAv7 5’-GAAACTCGCAATGGCTCATTAAATC-3’, 18sAv8

5’-TATTAGCTCTAGAATTACCACAGTTATCCA-3’). All samples were run in triplicate (Bio-Rad CFX96TM Real-Time System) and the mean of the two closet samples was used to calculate the threshold Ct value using the Bio-Rad CFX Maestro v1.1 software. Samples with a threshold Ct value higher than 35 were considered uninfected. The number of parasites were calculated as relative quantification values (RQ). RQ can be interpreted as the fold-amount of target gene (*Plasmodium* 18s rDNA) with respect to the amount of the reference gene (Bird18s rDNA) and are calculated as 2-^(C*t*18s *Plasmodium –* C*t*18s Bird)^. For convenience, RQ values were standardized by ×104 factor and log-transformed.

### Statistical analyses

Analyses were carried out using the R statistical package (v. 3.4.1). Data were analysed separately for each experiment and each experimental block.

Blood meal rate, blood meal size, infection prevalence, oocyst burden (where only individuals that developed ≥ 1 oocyst were included), quantity of parasites contained within the blood meal, which may depend on which bird mosquitoes fed on, were analysed fitting bird as a random factor into the models using *lmer, glmer* or *glmer.nb* (package: lme4, [77]) according to whether the errors were normally (haematin quantity, oocyst burden, quantity of parasites contained within the blood meal), binomially (blood meal rate, infection prevalence) or negative binomially distributed (oocyst burden). Mosquito batches (experience 1), mosquito biting order (experiment 2 and 3) and blood meal size (when it was not a response variable) were used as fixed factors. Parasitaemia and gametocytaemia of birds were analysed using *lmer* with bird fitted as a random factor into the models to account for temporal pseudo-replication. Times of day (5:45 and 9:00 p.m.) and bird group (exposed to mosquito bites or control) were used as fixed factors.

The different statistical models (maximal and minimal models) built to analyse the data are described in the supplementary material (**Table S1**). Maximal models, including all higher-order interactions, were simplified by sequentially eliminating non-significant terms and interactions to establish a minimal model [78]. The significance of the explanatory variables was established using a likelihood ratio test [79]. The significant Chi-square given in the text are for the minimal model, whereas non-significant values correspond to those obtained before the deletion of the variable from the model. *A posteriori* contrasts were carried out by aggregating factor levels together and by testing the fit of the simplified model using an LRT [78].

### Ethics statements

This study was approved by the Ethical Committee of the Vaud Canton veterinary authority, authorization number 1730.4.

### Data Accessibility

All data supporting the conclusions of this paper will be available on the Dryad website after acceptance.

## Supporting information captions

**Table S1: Description of statistical models presented in the main text**. “N” gives the number of mosquitoes or birds included in each analysis. “Maximal Model” includes the complete set of explanatory variables. “Minimal model” gives the model containing only the significant variables and their interactions. Square brackets indicate variables fitted as random factors. Curly brackets indicate the error structure used (n: normal errors, b: binomial errors, nb: negative binomial errors).

